# Evolution of vertical and oblique transmission under fluctuating selection

**DOI:** 10.1101/229179

**Authors:** Yoav Ram, Uri Liberman, Marcus W. Feldman

## Abstract

Vertical and oblique cultural transmission of a dichotomous phenotype is studied under constant, periodic cycling, and randomly fluctuating selection. Conditions are derived for the existence of a stable polymorphism in a periodically cycling selection regime. Under such a selection regime, the fate of a genetic modifier of the rate of vertical transmission depends on the length of the cycle and the strength of selection. In general, the evolutionarily stable rate of vertical transmission differs markedly from the rate that maximizes the geometric mean fitness of the population. The evolution of rules of transmission has dramatically different dynamics from the more frequently studied modifiers of recombination, mutation, or migration.

---

Cavalli-Sforza and Feldman (1) distinguished two forms of non-parental phenotypic transmission in the context of cultural evolution. Horizontal transmission occurs when a trait is passed between members of the same generation and is analogous to transmission of an infectious agent. Oblique transmission to offspring is from non-parental members of the parental generation. Evolution under either of these is expected to be more rapid than under purely vertical, i.e., parent-to-offspring, transmission (2, 3).

Oblique transmission occurs via some mechanism of social learning, which may include imitation or active teaching. There has been an interesting debate over the past thirty years concerning the conditions under which social learning would have an advantage over individual learning or vertical (including genetic) transmission. This debate is usually couched in terms of the mode and tempo of environmental fluctuations that would affect fitness and hence evolution (4–11). Mathematical analyses of models of competition between individual and social learning have generally shown that social learning has an advantage when the environment does not fluctuate too frequently. On the other hand, when the environmental changes are very frequent, individual learning is favored, while innate (genetic) determination of the trait does best when periods between environmental change are long on average.

In some situations, oblique transmission of biological material is possible. In bacteria, phenotypes might be determined by heritable mobile genetic elements such as phages (12), plasmids (13), integrons (14), and transposons (15). Similarly, some phenotypes are determined by genes that are commonly converted by uptake of foreign DNA, i.e., transformation (16). In these cases, inheritance of a phenotype may combine vertical transmission from the parent cell, and oblique transmission from other cells, even if the latter did not originally evolve for that purpose (17).

In some animals, transmission of microbes may occur during sharing or manipulation of food or other consumable resources during a social interaction. Although transmission of the microbiome in humans is likely to be mostly vertical (18), in other organisms there is multi-generational food sharing during which symbionts from the parental cohort may be transmitted obliquely to younger individuals (19). In such cases, fluctuations in the resource type or availability may have fitness effects that depend on features of the transmitted microbiome. This ecological perspective on community transmission is stressed by van Opstal and Bordenstein (20) who emphasize the “need to consider the relative roles of vertical and horizontal transmission of microbial communities.”

Another perspective on the evolutionary consequences of fluctuating environments (and, as a result, fluctuating selection) derives from the phenomenon of phenotypic switching (21–26) In these studies, mutation causes the organism to switch phenotypes (usually treated as haploid genotypes), and the problem has usually been couched in terms of the optimal rate of mutation in models where the phenotypic fitnesses fluctuate over time. These models did not include social learning, and the evolution was regarded as a mode of bet-hedging against future environmental change. Optimal (that is, evolutionarily stable) mutation rates depend on many features of the fluctuations, for example, degree of fitness symmetry, strength of selection, and variance in the period of fluctuation (24).

In a recent analysis of evolution under fluctuating selection, Xue and Leibler (27) allowed an organism to absorb information about the distribution of possible environments by learning the phenotypes of members of its parental lineage from previous generations. They describe this as “positive feedback that enhances the probability of the offspring to express the same phenotype as the parent.” In this formulation there was “reinforcement of the parent phenotype” in an offspring, such as might occur through epigenetic inheritance. Although their analysis was not couched in terms of oblique and vertical transmission, as defined by Cavalli-Sforza and Feldman (1), we have been stimulated by their analysis to develop a model in which oblique transmission, at a rate dependent on the trait frequency in the parental generation, occurs in addition to classical vertical transmission. We then ask how fluctuations in selection interact with the rate of oblique transmission to affect evolutionary dynamics and how the rate of oblique transmission itself might evolve.

In our formulation, both the parental phenotype and the distribution of phenotypes in the whole population contribute to an offspring’s phenotype. Using conventional modifier theory (28), we show that in a symmetric cyclic selection regime with cycles of periods 1 or 2, an allele reducing the rate of vertical transmission is expected to increase in frequency when rare and in so doing to increase the mean fitness of the population. However, for cycles of greater length or period asymmetry, interesting non-monotonicities emerge, both in the uninvadable rate of vertical transmission, and the rate that maximizes the geometric time average of the population mean fitness, which we will refer to as the “geometric mean fitness.” We develop the models in very large populations with cyclic selection and with random fitnesses and also in the case where drift occurs via sampling from generation to generation in a finite population.

## 1. The Model

Consider an infinite population whose members are characterized by their phenotype *ϕ*, which can be of two types *ϕ* = *A* or *ϕ* = *B*, with associated frequencies *x* and (1 − *x*), respectively. We follow the evolution of *x* over discrete non-overlapping generations. In each generation individuals are subject to selection where the fitnesses of *A* and *B* are *w_A_* and *w_B_*, respectively.

An offspring inherits its phenotype from its parent via *vertical transmission* with probability *ρ* and from a random individual in the parental population via *oblique transmission* with probability (1 − *ρ*). Therefore, given that the parent phenotype is *ϕ* and assuming *uni-parental inheritance* (29), the conditional probability that the phenotype *ϕ*′ of the offspring is *A* is

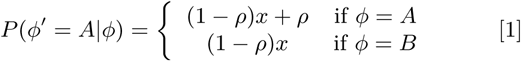

where *x* = *P*(*ϕ* = *A*) in the parent’s generation before selection.

Therefore the frequency *x*′ of phenotype *A* after one generation is given by the recursion equation

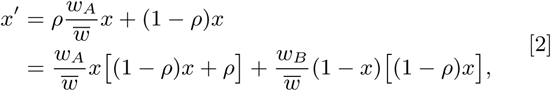

where *w̅* is the *mean fitness,* namely

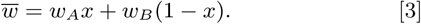

eq(2) can be rewritten as

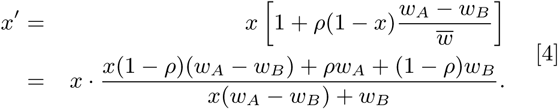

In what follows we explore the evolution of the recursion Eq. (4), namely the equilibria and their stability properties, in the cases of *constant environments* and *changing environments.*

### A. Constant Environment

When the environment is constant, the fitness parameters *w_A_* and *w_B_* do not change between generations, and we have the following result.

#### Result 1.

*If*0 *< ρ* ≤ 1 *and both w_A_* and *w_B_* are positive with *w_A_* ≠ *w_B_*, *then fixation in the phenotype A (B) is globally stable when w_A_* > *w_B_* (*w_A_* < *w_B_*).

#### Proof.

If we rewrite Eq. (4) as *x*′ = *x* · *f*(*x*), it can be seen that *f* (1) = 1, and for *ρ* > 0 and 0 < *x* < 1

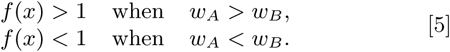

Hence, as *w_A_* > 0 and *w_B_* > 0, both fixations in *A* or in *B* (*x*\* = 1 for fixation in *A* and *x*\* =0 for fixation in *B*) are equilibrium points of Eq. (4). Moreover, if *x_t_* is the value of *x* at the *t*-th generation (*t* = 0,1, 2,…), from Eq. (4) and Eq. (5) we have for any 0 < *x*_0_ < 1 and all *t* = 0,1, 2,…,

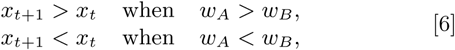

and since *x*\*=1 or *x*\* = 0 are the only equilibrium points,we have

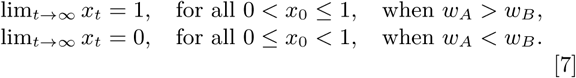

Therefore fixation of the favored phenotype is globally stable.

If *ρ* = 0 or *w_A_* = *w_B_* > 0, the phenotype frequencies do not change over time. With *extreme selection,* for example *w_A_* = 1 and *w_B_* = 0 (or *w_B_* = 1 and *w_A_* = 0), Eq. (4) reduces to *x*′ = (1 − *ρ*)*x* + *ρ*, and therefore *x_t_* = 1 − (1 − *ρ*)^*t*^ (1 − *x*_0_). Hence, for *ρ* > 0, *x_t_* → 1 as *t* → ∞ for all initial values 0 ≤ *x*_0_ ≤ 1, and the favored phenotype fixes.

### B. Periodically Changing Environment.

Suppose the environment is not constant but rather changes periodically, such that the favored phenotype changes after a fixed number of generations. Simple examples are *A*1*B*1 = *ABABAB*.‥, in which the favored phenotype switches every generation, or *A*2*B*1 = *AABAABAAB*‥., where every two generations in which selection favors *A* are followed by a single generation in which selection favors *B*. In general, *AkBl* denotes a selection regime in which the period is of (*k* + *l*) generations with *k* generations favoring phenotype *A* followed by *l* generations favoring *B*.

Let *W* be the fitness of the favored phenotype and *w* be that of the other phenotype where 0 < *w* < *W*. Rewrite Eq. (4) as *x*′ = *F_A_*(*x*) = *xf_A_*(*x*) when *A* is favored and *x*′ = *F_B_*(*x*) = *xf_B_*(*x*) when *B* is favored. Then

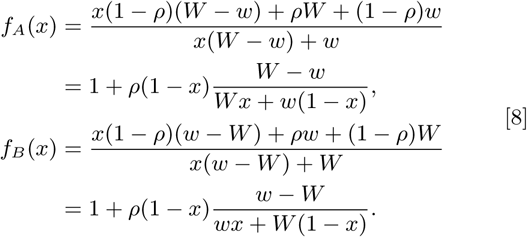

If *x_t_* denotes the frequency of the phenotype *A* at generation *t* starting with *x*0 initially, then as we are interested in the values of *x_t_* for *t* = *n*(*k* + *l*) with *n* = 0,1,‥. at the end of complete periods, we can write

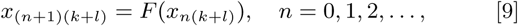

Where *F* is the composed function

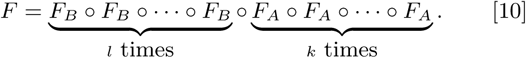

Clearly, since *F_A_* (0) = *F_B_* (0) = 0 and *F_A_* (1) = *F_B_* (1) = 1, both fixations in *A* or in *B* are equilibrium points. An interesting question is when these fixations are locally stable. We concentrate on *x*\* = 0, the fixation of the phenotype *B*.

As *x*′ = *F_A_*(*x*) = *xf_A_*(*x*)for *k* generations and *x*′ = *F_B_*(*x*) = *xf_B_*(*x*)for *l* generations, the linear approximation of *F*(*x*)“near” *x* = 0 is

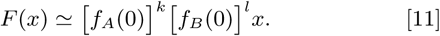

Hence the local stability of *x*^*^ = 0 is determined by the product [*f_A_*(0)]^*k*^[*f_B_*(0)]^*l*^; *x*\* = 0 is locally stable if this product is less than 1 and unstable if it is larger than 1.

From Eq. (8) we have

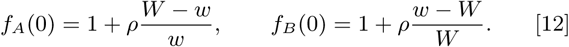

We start with the case *k* = *l*.

#### Result 2.

*If k* = *l and* 0 < *w < W with* 0 < *ρ* < 1, *fixation of B is unstable.*

*Proof.* The local stability of *x*\* = 0, the fixation of *B*, is determined by the product

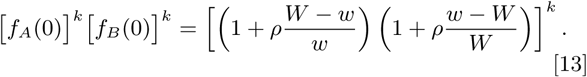

Observe that

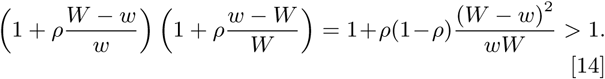

Since 0 < *ρ* < 1 and 0 < *w* < *W*, fixation on *B* is unstable.

#### Conclusions

1. Since *k* = *l* and the above result also holds when 0 < *W* < *w*, there is total symmetry between the two fixations in *A* and *B*, and fixation in *A* is also not stable. Thus neither phenotype can be lost, and there is a *protected polymorphism* (30).
2. For general *k*, *l*, the condition for local stability of fixation in *A* is

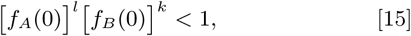

and that of *B* is

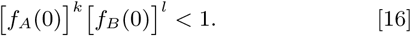 Therefore, following Result 2,

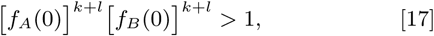

and it is impossible that both fixations are stable. Further, since by Eq. (12) *f_A_*(0) > 1 and 0 < *f_B_* (0) < 1 when 0 < *w* < *W*, by choosing *k* and *l* appropriately, fixation on *A* or fixation on *B* (but not both) can be stable. In addition, we can have both fixations unstable giving the following result.

##### Result 3.

*With* 0 < *ρ* < 1 and 0 < *w* < *W in the case of AkBl periodically changing environments, both fixations may be unstable, producing a protected polymorphism.*

##### Proof.

Let 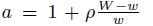 and 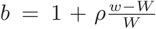 and our assumption entails *a* > 1 and 0 < *b* < 1. Following Eq. (11), fixation in *B* is not stable if *a^k^b^l^* > 1, and similarly fixation in *A* is unstable if *a^l^b^k^* > 1. Therefore both fixations are unstable if

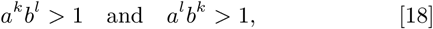

or equivalently if

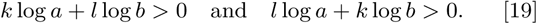

Now Eq. (19) holds if and only if

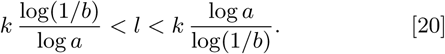

The inequalities of Eq. (20) are consistent if and only if log(1/*b*)< log *a*, i.e., *ab* > 1, which according to Eq. (14), is true.

The local stability properties of the two fixations depend only on the fact that in a cycle of (*k* + *l*) generations *A* is favored *k* times and *B* is favored *l* times, and not their order in the cycle. When neither fixation in *A* or *B* is stable, there is a protected polymorphism, and we expect to have one or more polymorphic equilibria. Supplementary Figure S1 illustrates the relationship between *k*, *l* and *ρ* that gives polymorphism of *A* and *B*, or fixation, for different values of *s* = *W* − *w*.

For the simple case of *A*1*B*1 periodically changing environment we have the following.

##### Result 4.

*In the case A*1*B*1 *with* 0 < *ρ* < 1 *and* 0 < *w* < *W the two fixations are unstable and there exists a unique stable polymorphism.*

##### Proof.

Let *x* be the initial frequency of *A* and *x*′ its frequency after one cycle of the *A*1*B*1 selection. Then *x*′ = *F_B_*(*F_A_*(*x*)) where by Eq. (8)

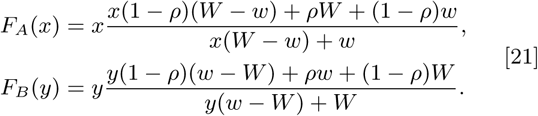

The equilibrium equation is *x* = *F_B_*(*F_A_*(*x*)), which reduces to a fourth degree polynomial equation in *x*. Since the two fixations in *B* and *A* are equilibria corresponding to the two solutions *x* = 0 and *x* = 1, the other equilibria correspond to solutions of a quadratic equation *Q*(*x*) = *α*_2_*x*^2^ + *α*_1_*x*+ *α*_o_ = 0 with

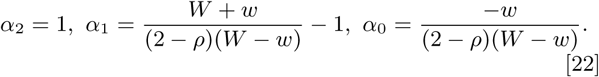

As 0 < *ρ* < 1 and 0 < *w* < *W* we have

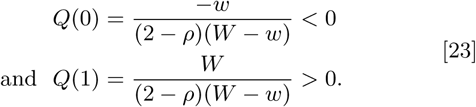

Also, as *α*_2_ = 1 and *α*_0_ < 0, the quadratic equation *Q*(*x*) = 0 has two real roots, one negative and one positive *x*\* satisfying 0 < *x*\* < 1. The latter determines a unique polymorphism. Let *H*(*x*) = *F_B_* (*F_A_*(*x*)). Then

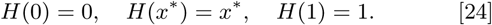

Also

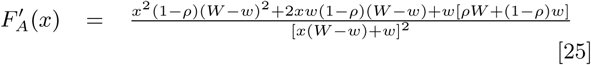

and

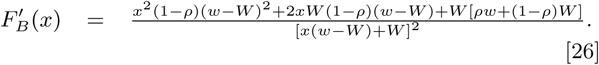

From our assumptions on *ρ*, *w*, and *W*, we have 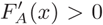 for 0 ≤ *x* ≤ 1. Observe that the numerator of 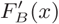 is linear in *ρ*; its value when *ρ* = 1 is *wW* > 0, and when *ρ* = 0 it is

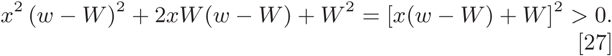

Hence 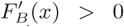 for all 0 ≤ *x* ≤ 1, and 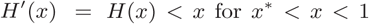 is positive when 0 ≤ *x* ≤ 1. Thus *H*(*x*) is monotone increasing for 0 ≤ *x* ≤ 1; *H*(*x*) > *x* for 0 < *x* < *x*\*,*H*(*x*) < *x* for *x*\* < *x* < 1. Starting from any initial value 0 < *x*_0_ < 1 we have *x_t_* → *x*\* as *t* → œ. Supplementary Figure S2, panels *A*, *C*, and *E*, illustrate examples of how the frequency of *A* changes over time in the *A*1*B*1 regime of cycling selection.

For more general cyclic fitness regimes, the polynomial that gives the equilibria is of higher order, and it is conceivable that more than one stable polymorphism could exist for given values of *ρ*, *W*, and *w*. We have been able to show that when neither fixation in *A* or *B* is stable, in the *AkBk* case this cannot occur. In fact we have the following.

##### Result 5.

*In the AkBk selection regimes, if the fixations in A and B are locally unstable, a single stable polymorphic equilibrium exists.*

The proof of Result 5 is in Supplementary Material SP1. Supplementary Figure S3A shows the stable equilibrium frequencies *x*\* as a function of *ρ*, *W*, and *w* in the *A*1*B*1 regime. For *AkBk* selection regimes from *k* = 1 to *k* = 40, Supplementary Figure S4 illustrates the convergence to a single stable polymorphism.

We have not been able to prove that for selection regimes *AkBl* with *l* ≠ *k* there is a single stable polymorphic equilibrium when the two fixations are unstable. However, the numerical examples in Supplementary Figure S1 (for *AkBl*) and in Figure 1 and Supplementary Figure S5 for the special case *A*1*B*2 all exhibit a single stable polymorphic equilibrium when fixations in *A* and *B* are unstable. These numerical results suggest that for *W* > *w* > 0 and 0 < *ρ* < 1 the high order polynomial that gives the equilibria has only a single root corresponding to a globally stable polymorphism. Supplementary Figure S6 shows that this is the case for the *A*3*B*10 regime.

**Fig. 1.**
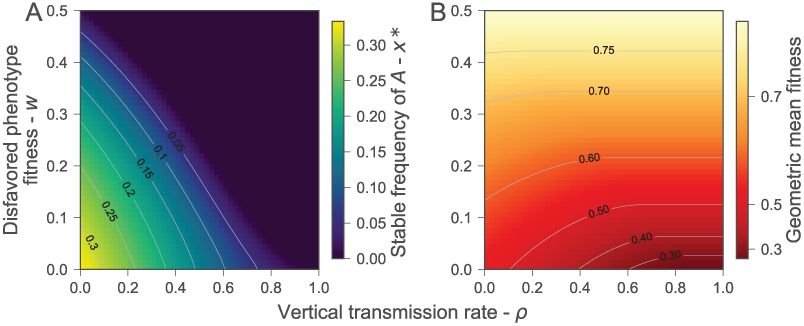
Stable frequency of phenotype *A* and geometric mean fitness in selection regime *A*1*B*2 as a function of the vertical transmission rate *ρ* and the fitness of the disfavored phenotype *w*. (**A**) Stable frequency of phenotype *A* at the end of each three generations cycle. (**B**) Geometric average of the stable population mean fitness over the three generations cycle: 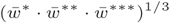. Gray contour lines join *ρ* and *w* combinations that result in the same stable value. In all cases, *W*=1.

### C. Randomly Changing Environment.

We now consider the case where the environment changes according to a stochastic process. Without loss of generality, assume that the fitness parameters at generation *t* (*t* = 0,1, 2,≠.) are 1 + *s_t_* for phenotype *A* and 1 for phenotype *B*, where the random variables *s_t_* for *t* = 0,1, 2… are independent and identically distributed. Also we assume that there are positive constants *C* and *D* such that *P*(−1 + *C* < *s_t_* < D) = 1.

Corresponding to Eq. (4), with *w_A_* = 1 + *s_t_*, *w_B_* = 1 the recursion equation is

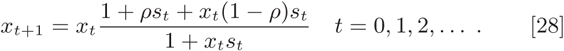

As {*x_t_*} for *t* = 0,1, 2,≠. is a sequence of random variables, the notion of stability of the two fixation states needs clarification. Following Karlin and Lieberman (31) and Karlin and Liberman (32) we make the following definition.

#### Definition

“stochastic local stability”. A constant equilibrium state *x*\* is said to be *stochastically locally stable* if for any *ε* > 0 there exists a *δ* > 0 such that |*x*_0_ − *x*\*| < *δ* implies

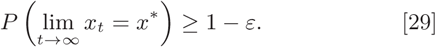

Thus stochastic local stability holds for *x*\* provided for any initial *x*_0_ sufficiently near *x*\* the process *x_t_* converges to *x*\* with high probability.

In our case there are two constant equilibria *x*^*^ = 0 and *x*^*^ = 1 corresponding to fixation in *B* and *A*, respectively. We can characterize the stochastic local stability of these fixations as follows.

##### Result 6.

*Suppose E* [log(1 + *ρs_t_*)] > 0. *Then x*\* = 0, *the fixation of phenotype B, is not stochastically locally stable. In fact P* (lim_*t*→∞_ *x_t_* = 0) = 0.

##### Result 7.

*Suppose E*[log(1 + *ρs_t_*)] < 0. *Then x*\* = 0, *the fixation of phenotype B, is stochastically locally stable. In particular, if E*(*s_t_*) ≤ 0, *x*\* =0 *is stochastically locally stable.*

The proofs of Results 6 and 7 can be found in Supplementary Materials SP2 and SP3.

Using the general notation for the fitness parameters *w_A_* and *w_B_*, the stochastic local stability of the *B* fixation is determined by the sign of 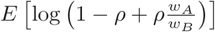and that of the *A* fixation by the sign of 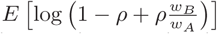. For example, if the sign of the first is negative, fixation in *B* is stochastically locally stable, and when it is positive, with probability one convergence to fixation in phenotype *B* does not occur. It is also true that if *E*(*w_A_*/*w_B_*) ≤ 1, then fixation of *B* is stochastically locally stable. Following Eq. (14), for all realizations of *w_A_* and *w_B_*,

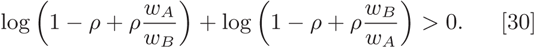

Therefore, as in the case of periodically changing environments *AkBl*, it is impossible that both fixations are simultaneously stochastically locally stable. It is possible, however, that neither fixation is stochastically locally stable, in which case we expect the population to converge to a polymorphic distribution. Figure 2 illustrates how the properties of *s_t_* in Eq. (28) affect the frequency of phenotype *A*, and in particular the stochastic local stability of fixation in phenotype *B*. Figure 3 shows the dynamics of the frequency of *A* in a case where *w_A_* and *w_B_* are identically distributed and independent; in this case the expectation of the stationary distribution is 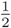, and its variance increases as *ρ* increases.

**Fig. 2.**
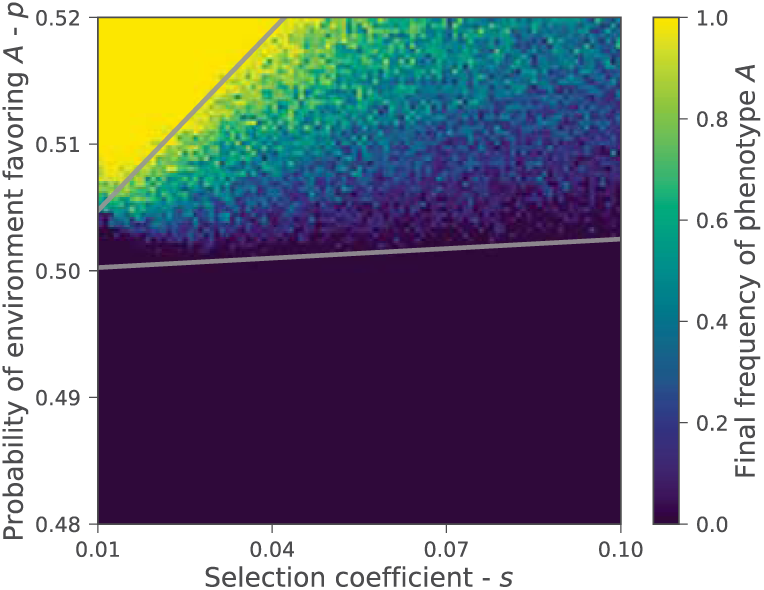
Stochastic local stability. The figure shows the frequency of phenotype *A* after 10^6^ generations in a very large population evolving in a stochastic environment (eq. 28). The fitnesses of phenotypes *A* and *B* are *w_A_* = 1 + *s_t_* and *w_B_* = 1, where *s_t_* is *s* with probability p and -*s* with probability 1 − *p*. The gray lines mark combinations of *p* and *s* for which 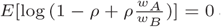 and 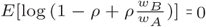. According to Result 6, between these lines fixation of either phenotype is not stochastically locally stable, and we expect a stationary polymorphism between the lines. Here, initial frequency of *A* is *x*_0_ = 1/10, 000 and the vertical transmission rate is *ρ* = 0.1.

**Fig. 3.**
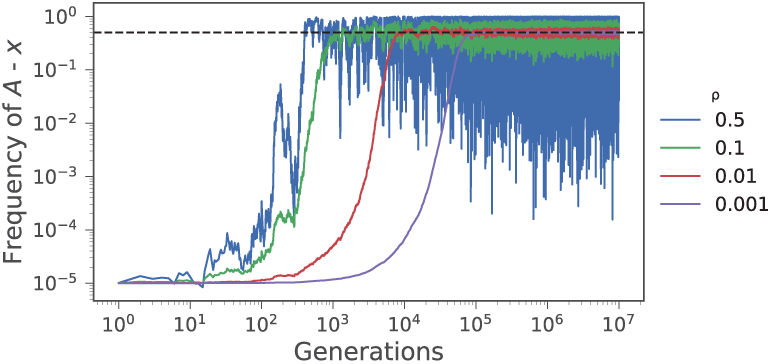
Effect of vertical transmission rate *ρ* on phenotype polymorphism in a randomly changing environment. Dynamics of the frequency of phenotype *A* over time starting at *x*_0_ = 10^−5^ when the fitnesses of phenotypes *A* and *B* are identically and independently distributed random variables. As the vertical transmission rate *ρ* increases from 0.001 to 0.5, the frequency reaches a polymorphic distribution with *E*(*x_t_*) → 0.5 faster, but the variance also increases. The fitnesses of phenotypes *A* and *B*, *w_A_* and *w_B_*, are both exponential random variables with expected value 2.

## 2. Evolutionary Stability of Oblique Transmission

An interesting question concerns the evolution of oblique transmission itself. For example, is there an evolutionarily stable rate of oblique transmission? To answer this question we use a modifier model, for which we suppose that the vertical transmission rate is controlled by a genetic locus with two possible alleles *m* and *M*. Let the vertical transmission rates determined by *m* and *M* be *ρ* and *P*, respectively. Thus there are four pheno-genotypes *mA*, *mB*, *MA*, *MB* whose frequencies at a given generation are denoted by *x*_1_, *x*_2_, *x*_3_, *x*_4_, respectively. As the fitnesses are determined by the two phenotypes *A* and *B*, and the modifier locus is selectively neutral, we have the following table.

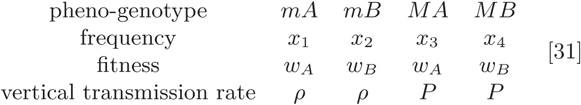

Following the rationale leading to equation [2], the next generation pheno-genotype frequencies 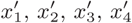 are

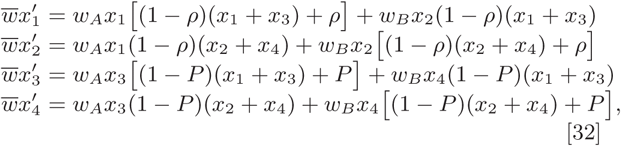

with *w̅,* the mean fitness, given by

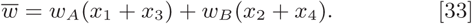

Note that under these assumptions the *M*/*m* locus and the *A*/*B* phenotypic dichotomy do not undergo anything analogous to recombination, which might be introduced if *A*/*B* phenotypes were viewed as haploid genetic variants.

Starting with a stable equilibrium where only the *m* allele is present, we check its *external stability* (28, 33) to invasion by allele *M*. Constant environment always leads to fixation of the favored type, independent of *ρ*. We therefore assume changing environments, and, in particular, the simple case of the *A*1*B*1 cycling environment, where a unique stable polymorphism exists and depends on *ρ* (but see Supplementary Material SP6 for computational analysis of the general *AkBl* case). Specifically, from Eq. (32) with *w_A_* = *W*, *w_B_* = *w* in the first generation and *w_A_* = *w*, *w_B_* = *W* in the second generation, after two generations we have

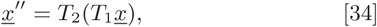

where 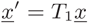 is given by Eq. (32) with *w_A_* = *W*, *w_B_* = *w*, and 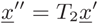 is given by [32] with *w_A_* = *w*, *w_B_* = *W*. Here 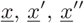 are the frequency vectors.

For the *A*1*B*1 case, when only the *m* allele is present with associated rate *ρ,* 0 < *ρ* < 1, and 0 < *w* < *W*, a unique stable equilibrium 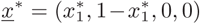 exists. 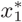 is the only positive root of the quadratic equation *Q*(*x*) = *α*_2_*x*^2^ + *α*_1_*x* + *α*_0_ = 0 with *α*_2_,α_1_,*α*_0_ specified in Eq. (22). Solving *Q*(*x*) = 0 gives

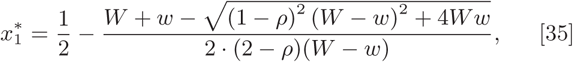

and it can be seen that

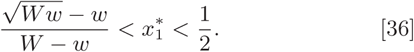

The external stability of 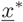 to the introduction of the modifier allele *M* with rate *P* is determined by the linear approximation matrix **L** = **L**_2_ · **L**_1_ near *x*\*, which is derived from Eq. (32) and given by

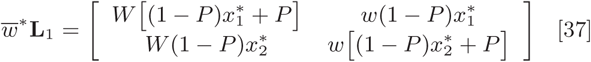

and

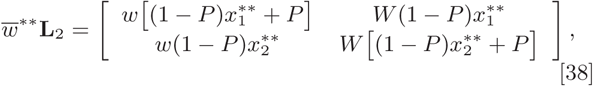

where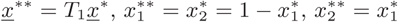and

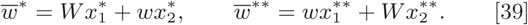

Due to the symmetry between the two phenotypes *A* and *B* in the *A*1*B*1 case, we have 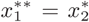 and 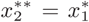 so that 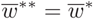 and in fact

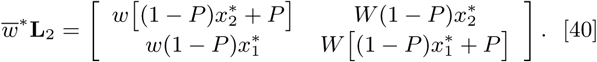

Note that as 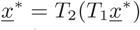 with 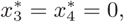, from Eq. (37) and Eq. (38) with *P* = *ρ* we have

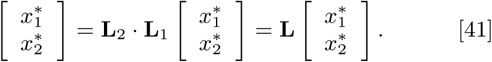

Hence when *P* = *ρ* one of the eigenvalues of **L** is 1.

In general **L** = **L**_2_ · **L**_1_, and using Eq. (37) and Eq. (40) we have

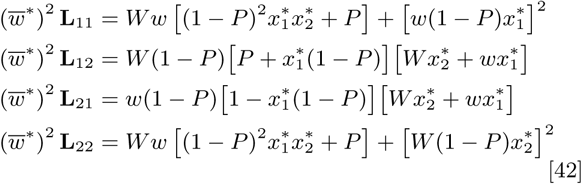

The external stability of 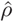 is determined by the eigenvalues of **L**, namely the roots of its characteristic polynomial *R*(*λ*) = det(**L** − *λ***I**), with **I** the 2 × 2 identity matrix. From Eq. (42), *R*(*λ*) = *a*_2_*λ*^2^ + *a*_1_*λ* + *a*_0_, where

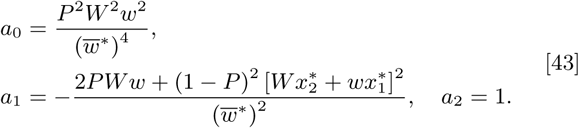

We then have the following result, which is proved in Supplementary Material SP4.

### Result 8.

**L** *has two positive eigenvalues and*

i. *when P* > *ρ the two eigenvalues are less than* 1,
ii. *when P* < *ρ the largest eigenvalue is larger than* 1,
iii. *when P* = *ρ*, *the largest eigenvalue is* 1.

We conclude that in the *A*1*B*1 selection regime an allele *m* producing vertical transmission rate *ρ* is stable to the introduction of a modifier allele *M* with associated rate *P* if *P* > *ρ*, and it is unstable if *P* < *ρ*. Thus in this case, evolution tends to reduce vertical transmission, and hence increase the rate of oblique transmission, and there is a *reduction principle* for the rate of vertical transmission (28, 33). The evolutionary dynamics of the reduction in *ρ* under the *A*1*B*1 cycling regime are shown in Figure 4, which also illustrates the change in phenotype frequencies over time.

**Fig. 4.**
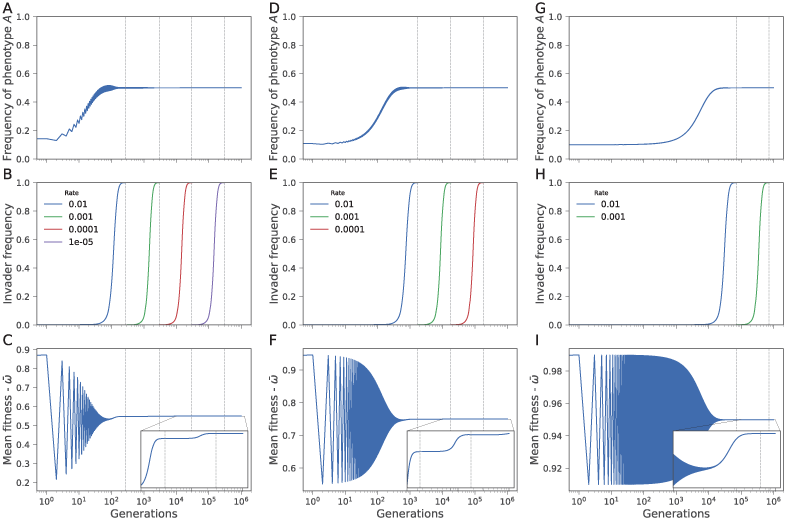
Consecutive fixation of modifiers that reduce the vertical transmission rate in selection regime *A*1*B*1. The figure shows results of numerical simulations of evolution with two modifier alleles (Eq. (32)). When a modifier allele fixes (frequency>99.9%), a new modifier allele is introduced with a vertical transmission rate one order of magnitude lower (vertical dashed lines). (**A,D,G**) The frequency of phenotype A in the population over time. (**B,E,H**) The frequency of the invading modifier allele over time. (**C,F,I**) The population geometric mean fitness over time; insets zoom in to show that the mean fitness decreases slightly with each invasion. Parameters: vertical transmission rate of the initial resident modifier allele, *ρ*_0_ = 0.1; fitness values: *W* = 1; *w* = 0.1 (**A-C**), 0.5 (**D-F**), and 0.9 (**G-I**). The x-axis is on a log-scale, as each sequential invasion takes an order of magnitude longer to complete.

In the case of identically distributed random fitnesses *w_A_* and *w_B_*, Figure 5 shows an example of the success of modifiers that reduce *ρ*. We have not, however, been able to prove that there is a reduction principle for this class of fluctuating fitnesses.

**Fig. 5.**
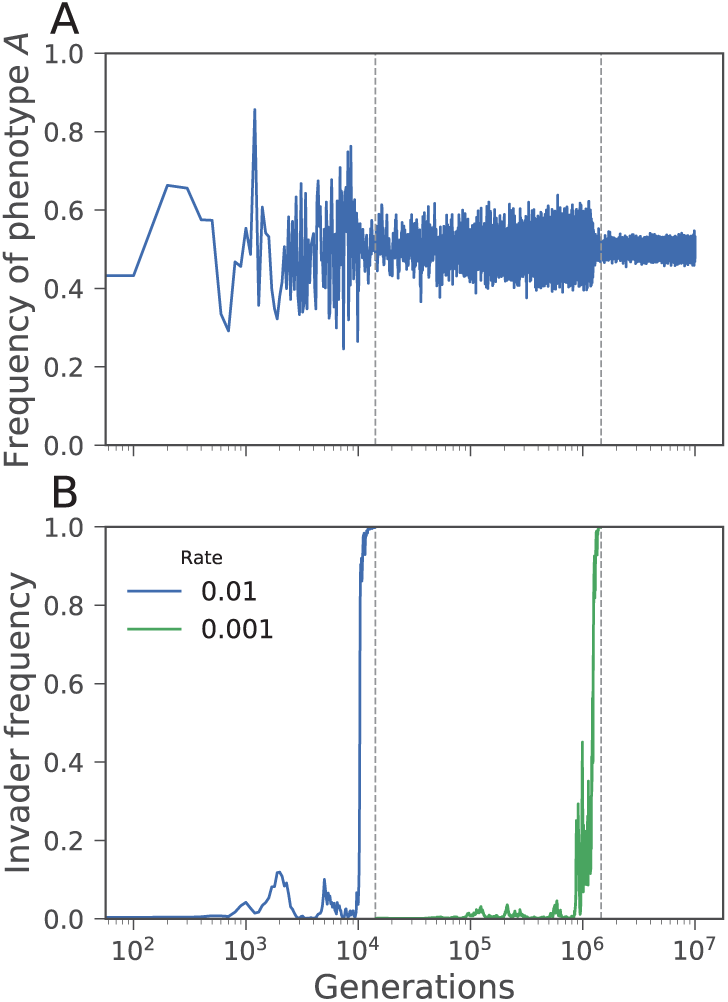
Consecutive fixation of modifiers that reduce the vertical transmission rate *ρ*under symmetric randomly changing selection. The figure shows results of numerical simulations of evolution with two modifier alleles (Eq. (32)). When a modifier allele fixes (frequency>99.9%), a new modifier allele is introduced with a vertical transmission rate one order of magnitude lower (vertical dashed lines). (**A**) The frequency of phenotype *A* in the population over time. (**B**) The frequency of the invading modifier allele over time. Parameters: vertical transmission rate of the initial resident modifier allele is *ρ*_0_ = 0.1 and the ratio of fitness values is *w_A_*/*w_B_* = 10 with probability 0.5 and *w_A_*/*w_B_* =0.1 also with probability 0.5. The x-axis is on a log-scale, as each sequential invasion takes an order of magnitude longer to complete.

Values of *ρ*\* for some *AkBl* examples (see Supplementary Material SP7 and Figure S10 for analytical details) are recorded in Table 1 for different values of *w* relative to *W* = 1. Interestingly, for the A1B2 regime, with *w* = 0.1, the evolutionarily stable value of *ρ* is substantially greater than zero, as it is in the *A*3*B*10 and *A*5*B*30 regimes. *AkBk* results are plotted in Figure 6B. In the *A*2*B*2 regime, *ρ*\* =0 and there is reduction of vertical transmission for all selection values tested. However, for *AkBk* regimes with *k* > 2, we find *ρ*\* ≠ 0, and depending on *w*, *ρ*\* can be as high as 0.95. In Table 1, blank values for *ρ*\* indicate that our method was numerically unstable and that a precise value for *ρ*\* could not be obtained. This is why, in Figure 6B, no *ρ*\* points are shown for *AkBk* with *k* > 19. In Table 1, the word “fixation” indicates that fixation of *B* occurs, at which point there can be no effect of modification of *ρ*; *ρ*\* cannot be calculated in such cases.

**Fig. 6.**
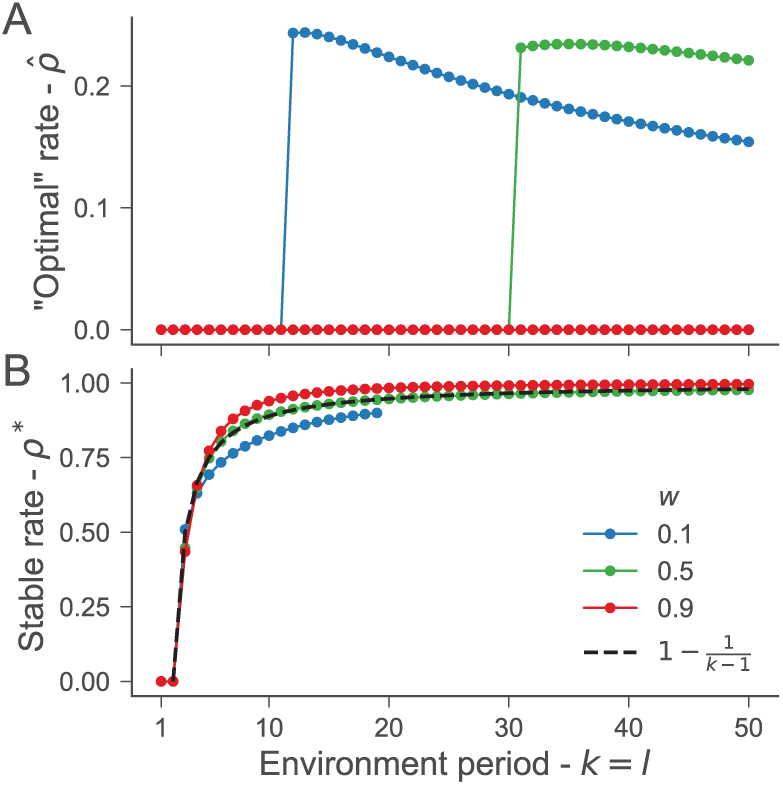
Fitness “optimal” and evolutionary stable vertical transmission rate in *AkBk* selection regime. (**A**) The vertical transmission rate 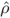 that maximized the geometric average of the population mean fitness is zero (complete oblique transmission) when selection cycles quickly between favoring phenotype *A* and *B*, and then abruptly transitions to ≈ 0.2, followed by a slow decrease (see Fig. S5 and S6 for details on the abrupt transition). (**B**) The evolutionary stable rate *ρ*\*, which cannot be invaded by modifiers with either higher or lower vertical transmission rate *P*, rapidly increases from zero when selection cycles are short (*k* = 1 or 2) to roughly 1 when selection cycles are longer. The dashed line shows 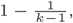, which fit the values for *w* = 0.5(Carja et al. 2011). The values for *w* = 0.1 (blue) could not be calculated for *k* > 19 due to numerical instability when selection is strong and the duration between selection fluctuations is long. In all cases, *W* =1. See *SupplementalMaterial SP6* for details on how we calculated the stable rate.

## 3. Geometric Mean Fitness and Rate of Vertical Transmission

Under fluctuating selection, the geometric mean fitness of genotypes has been shown to determine their evolutionary dynamics (8, 31, 34). For the evolution of mutation rates that are controlled by genetic modifiers, the stable mutation rate and the mutation rate that maximizes the geometric mean fitness of the population appear to be the same when the period of environmental fluctuation is low enough (25). We can ask the same question here: is the stable rate *ρ*\* the same as the rate 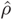 that maximizes the equilibrium value of the geometric mean fitness under fluctuating selection. First consider the *A*1*B*1 selection regime. We have the following result.

### Result 9.

*If W* > *w and* 0 ≤ *ρ* ≤ 1, *then the mean fitness at the stable equilibrium in the A*1*B*1 *environment is a decreasing function of ρ.*

### Proof.

In *A*1*B*1 the stable frequency of phenotype *A* is by Eq. (35)

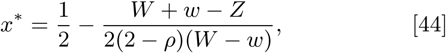

where 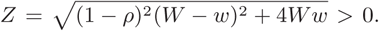. The mean fitness at the stable equilibrium is 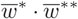, and as 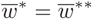by Eq. (39), this allows us to reduce the problem to properties of 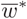. Now since 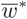 is an increasing linear function of*x*\*:

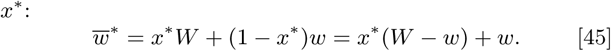

Thus 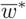 is decreasing in *ρ* if *dx*\* /*dρ* is negative. Using Eq. (44),

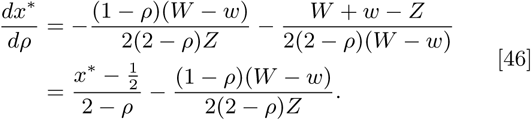

From Eq. (36) 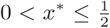 and therefore *dx*\* /*dρ* < 0, which completes the proof.

Figure 4 illustrates the decrease over time of the geometric mean fitness with increasing *ρ* at a polymorphic equilibrium in the *A*1*B*1 regime. The values of 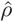 and *ρ*\* are the same in *A*1*B*1 and *A*2*B*2 regimes, namely both are zero. Figure 1B shows the geometric mean fitness in the *A*1*B*2 regime, and we see that for small values of *w*, this mean fitness decreases as *ρ* increases. At *w* = 0.1, Table 1 shows that 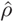 is 0.00065, whereas *ρ*\* = 0.821. In all *AkBk* regimes that we tested with *w* = 0.9, the value of 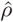 was also zero, substantially different from the values of *ρ*\*, as shown in Figures 6A and 6B. Also in Figure 6A, we see that with *w* = 0.1, 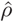 changes from zero to positive in the *AkBk* regimes with *k* ≥ 12, while with *w* = 0.5, the change occurs at *k* = 31. In Figure 6, with *w* = 0.1, 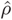 is between 0.15 and 0.24 for 12 ≤ *k* ≤ 50, while with *w* = 0.5, 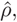 exceeds 0.2 for 31 ≤ *k* ≤ 50. More details on the mismatch between *ρ*\*, which cannot be invaded, and 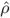, which maximizes geometric mean fitness, are given in Table 1, Supplementary Figures S7 and S8, and Supplemental Material SP7.

**Table 1.** Values of *ρ*\* (stable ρ)^(a)^ and 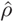 (“optimal” *ρ*)^(b)^

| $k$ | $l$ | $w^\dagger$ | $\hat{\rho}$ | $\rho^*$ |
| --- | --- | --- | --- | --- |
| 1 | 1 | 0.1 | 0.0000000 | 0.0000000 |
| 1 | 1 | 0.5 | 0.0000000 | 0.0000000 |
| 1 | 1 | 0.9 | 0.0000000 | 0.0000000 |
| 1 | 2 | 0.1 | 0.00065 | 0.821 |
| 1 | 2 | 0.5 | fixation | fixation |
| 1 | 2 | 0.9 | fixation | fixation |
| 2 | 2 | 0.1 | 0.0000000 | 0.0000000 |
| 2 | 2 | 0.5 | 0.0000000 | 0.0000000 |
| 2 | 2 | 0.9 | 0.0000000 | 0.0000000 |
| 3 | 10 | 0.1 | 0.00031 | 0.619 |
| 3 | 10 | 0.5 | fixation | fixation |
| 3 | 10 | 0.9 | fixation | fixation |
| 5 | 30 | 0.1 | 0.00027 | 0.6503 |
| 5 | 30 | 0.5 | fixation | fixation |
| 5 | 30 | 0.9 | fixation | fixation |
| 12 | 12 | 0.1 | 0.24347 | 0.84924 |
| 12 | 12 | 0.5 | 0.0000000 | 0.91209 |
| 12 | 12 | 0.9 | 0.0000000 | 0.95686 |
| 20 | 20 | 0.1 | 0.223925 |  |
| 20 | 20 | 0.5 | 0.0000000 | 0.94643 |
| 20 | 20 | 0.9 | 0.0000000 | 0.98304 |
| 30 | 30 | 0.1 | 0.1932801 |  |
| 30 | 30 | 0.5 | 0.0000000 | 0.96331 |
| 30 | 30 | 0.9 | 0.0000000 | 0.0000000 |
| 50 | 50 | 0.1 | 0.15419 |  |
| 50 | 50 | 0.5 | 0.22107 | 0.9768 |
| 50 | 50 | 0.9 | 0.0000000 | 0.99581 |

<sup>†</sup>Note: $W = 1$ .
(a): $\rho^*$ is the uninvadable value of the vertical transmission rate.
(b): $\hat{\rho}$ is the geometric mean value of $\rho$ at the stable equilibrium of the $AkB1$ cycle.

## 4. Finite Population Size

In order to include the effect of random drift due to finite population in the above deterministic model, we use the Wright-evolutionary stable vertical transmissionFisher model. Let *X_t_* denote the number of individuals with phenotype *A* in a population of fixed size *N* at the *t*-th generation, and suppose *X_t_* = *Nx.* Also, let *x*′ represent the frequency of the phenotype *A* in the infinite population model in the next generation, namely (see Eq. (2)),

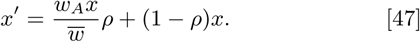

Then, according to the Wright-Fisher model (35), *X_*t*+1_,* the number of individuals of phenotype *A* at generation (*t* + 1), is determined by the probability

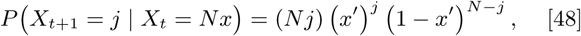

for *j* = 0,1, 2,….,*N*. Thus the fluctuations in the numbers of phenotypes *A* and *B* in the population of size *N* are generated by the Wright-Fisher Markov chain where, given that *X_t_* = *Nx*, *X*_*t*+1_ has a binomial distribution with parameters (*N*,*x*′).

This Markov chain has two absorbing states, *X_t_* = *N* and *X_t_* = 0, corresponding to the two fixations in *A* and *B*,respectively, and we are interested in the fixation probabilities and the time to fixation of these two absorbing states as functions of the initial frequency *x* and also of *ρ*, *w_A_*, and *W_B_*. To these ends we use a diffusion approximation to the process {*X_t_*}. Specifically, we compute the mean *μ*(*x*) and the variance *σ*^2^(*x*) of the change in one generation in the frequency of phenotype *A* given that at the beginning of the generation *X_t_* = *Nx*.

To compute *μ*(*x*), observe that by Eq. (47)

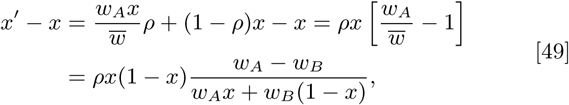

since *w̅* = *w_A_x* + *w_B_*(1 − *x*). For the diffusion approximation, it is essential that the differential selection does not have a large effect per individual in each time period 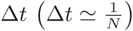. That is, we assume

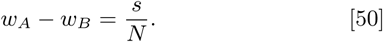

Then

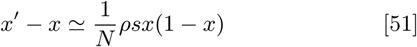

up to terms of order small than 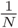. Since one generation corresponds to 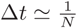, we conclude that

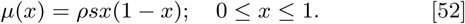

In the same way, we can compute *σ*^2^(*x*), which is *x*(1 − *x*). These values of *μ*(*x*) and *σ*^2^*x*) allow us to compute *u*(*x*),Namely

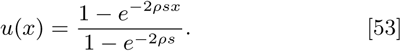

The expected time to fixation in *A* starting from a initial frequency of *x* is is given by

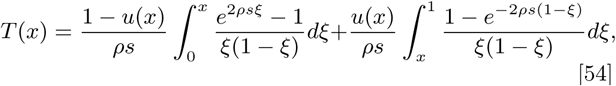

where *u*(*x*) is given in Eq. (53), and in generations, *T*(*x*) is multiplied by *N*. Unfortunately the integrals in Eq. (54) cannot be done in closed form unless *ρs* = 0, in which case *u*(*x*) = *x* and *T*(*x*) = −2*x*ln*x* − 2(1 − *x*) ln(1 − *x*) as Ewens points out on p. 160 (35), and only numerical computation of *T*(*x*) is possible for specified values of *x*, *ρ*, and *s*.

It is important to note that the fixation probability *u*(*x*) is a monotone increasing function of *ρ* when *s* > 0. In fact we have the following result.

### Result 10.

*When s* > 0 *so that the phenotype A is favored, the fixation probability u*(*x*) *is monotone increasing in ρ.*

The proof of Result 10 is in Supplementary Material SP5. Supplementary Figure S9 compares the fixation probability and time to fixation derived numerically from simulating the Wright-Fisher Markov chain with the diffusion-derived values of *u*(*x*) and *T*(*x*). The fit is seen to be very good. Note that when *N* is large, the Wright-Fisher model exhibits persistent fluctuation around the deterministic expectation, as shown by the orange diagram in Supplemental Figure S2.

We can also develop a diffusion approximation for the case of a cycling environment. Suppose that selection changes in cycles of length *n* such that within the cycle the fitness parameters are 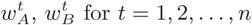 for *t* =1, 2,…,*n*. Also let

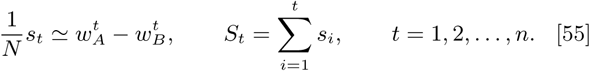

Following Karlin and Levikson (36) we have the following result.

### Result 11.

*The mean μx*) *and variance σ*^2^(*x*) *of the change in the frequency of A in one generation for the diffusion approximation in the case of a cycling environment AkBl, where k* + *l* = *n, are*

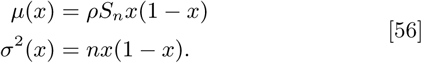

The proof of Result 11, based on induction on *n*, is given in Supplemental Materials SP6.

Using the moments in Eq. (56), the fixation probability *u*(*x*) and the expected time *T*(*x*) to fixation from an initial frequency of *x* can be computed where *s* is replaced by *s_n_*/*n*.We find

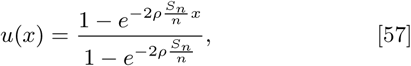

and *T*(*x*) can be computed similarly.

In the case of the *AkBl* cycling environment, we write *n* = *k* + *l*, and if *w_A_* = *W*, *w_B_* = *w* for *k* generations, and *w_A_* = *w*, *w_B_* = *W* for *l* generations, we have

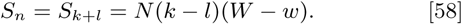

Figure 7 shows an example of how (*k*− *l*), *ρ*, and (*W*−*w*), which enter the formula for *u*(*x*) in Eq. (57), interact to affect fixation probabilities. More examples are illustrated in Supplementary Figure S11.

**Fig. 7.**
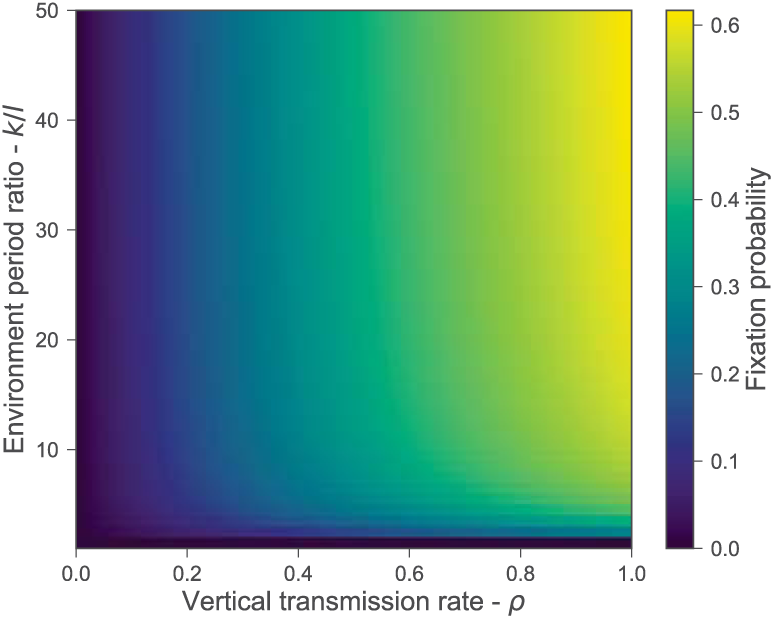
Fixation in a finite population with different ratios of selection periods 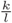. Fixation probability of phenotype *A* when starting with a single copy in a population of size 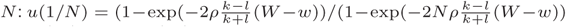 (see Eq. (57) and Eq. (58)). *k* and *l* are the number of generations in which phenotypes *A* and *B,* respectively, are favored by selection. Here, fitness of the favored phenotype is *W* = 1, fitness of the unfavored phenotype is *w* = 0.5, and the population size is *N* = 10, 000. See Figure S11 for additional examples.

## Discussion

Non-chromosomal modes of phenotypic transmission are receiving increasing attention (37–39), especially with respect to their potential role in adaptation and maintenance of diversity (40). Here we have focused on a dichotomous phenotype transmitted through a combination of parental and non-parental transmission. In addition to the roles that these transmission modes play in the dynamics of phenotypic diversity in large and small populations, we have also investigated a genetic model for the evolution of the transmission mode itself.

Our model differs markedly from that of Xue and Leibler (27), who took the individual phenotypic distribution (i.e., the probability that an individual develops one of a set of phenotypes) to be the inherited trait. In our model, the transmitted trait is the phenotype itself. Thus, with two phenotypic states *A* and *B*, we track the frequency *x* of *A,* whereas Xue and Leibler focus on the dynamics of the per-individual probability *π*_A_ of learning the phenotype *A*. One interpretation of our model is as a mean-value approximation to the model of Xue and Leibler, where *x*, the state in our model, is the average of the population distribution of individual phenotype probabilities.

In a constant environment, the higher the vertical transmission rate *ρ*, the faster is the approach to fixation of the favored phenotype: *A* if *w_A_* > *w_B_* or *B* if *w_B_* > *w_A_*. Here 1 − *ρ*, the oblique transmission rate, represents the added chance that an offspring becomes *A* by learning from the parent’s population after learning from its parents who have undergone selection (eq. 2). This simple phenotypic model does not allow a polymorphism to be achieved in a constant environment, but with more oblique transmission, approach to fixation is retarded.

With fluctuating environments, the dynamics of the phenotype frequencies are, in general, much more complicated. In particular, with deterministically cycling symmetric fitness values (the *AkBl* model), it is impossible for fixation in *A* and *B* to both be stable. If *k* = *l*, for example, neither fixation is stable, and there is a single stable polymorphic equilibrium (with phenotypes *A* and *B* present; see Result 5). In the *A*1*B*1 case this polymorphism is globally stable. In the *AkBl* case, bounds on *l*/*k* that determine the instability of both fixations, and hence the protection of polymorphism are given by the inequalities of Eq. (20), which depend on both the fitness differences and the rate *ρ* of vertical transmission. We conjecture that with *k* ≠ *l* there is a unique stable polymorphism when both fixations are unstable.

In deterministic one-locus, two-allele diploid population genetic models with cycling fitness regimes, Haldane and Jayakar (34) first showed the relevance of the geometric mean of genotypic fitnesses (cf. Eq. (16) and Eq. (17)) for the maintenance (or loss) of polymorphism. However, with equal homozygote fitnesses, which alternated in strength as a two-generation cycle (cf. *A*1*B*1), Karlin and Liberman (32) extended the Haldane-Jayakar results and found conditions under which both allelic fixations and polymorphic equilibrium could all be stable, with the evolution depending on initial allele frequencies, as well as the homozygote fitness differences between alternate generations. Our haploid model appears not to produce such dependence on the initial conditions.

When the fitnesses *w_A_* and *w_B_* are treated as random variables, rather than varying cyclically, stochastic local stability is the appropriate analog to local stability in the case of cyclic fitness variation. While fixations in phenotypes *A* and *B* cannot both be stochastically stable in this case, both may be unstable and a polymorphic distribution may result. The variance of this distribution is greater for larger values of *ρ*. This is because the stochastic local stability conditions involve *E*{log[1 − *ρ* + *ρ*(*w_A_*/*w_B_*)]} and the effect of the variance of (*w_A_*/*w_B_*) will clearly increase as *ρ* increases. In the finite population case, a greater level of vertical transmission makes selection more effective, increasing the probability *u*(*x*) of fixation and reducing the expected time to fixation.

We have shown that in the *A*1*B*1 case, the rate of vertical transmission tends to decrease when it is under the control of a genetic modifier. From numerical iteration, it appears that this is also true in the random selection case, when the fitnesses of *A* and *B* are identically distributed and independent between generations. However, for *AkBl* selection regimes more complicated than *A*1*B*1, evolution of a modifier of vertical transmission is not straightforward. While reduction of *ρ* occurs in the *A*2*B*2 regime, the uninvadable value *ρ*\* is not zero for all the fitness values explored in *AkBk* regimes with *k* > 2 (Figure 6B, Table 1). In fact, *ρ*\* increases sharply as *k* increases beyond *k* = 2. This is an unusual scenario for genetic modifiers, although it must be noted that a modifier of *ρ* is not neutral; it affects primary selection, while neutral modifiers of recombination, mutation, and migration affect induced or secondary selection.

The dependence of the modifier dynamics on the strength of selection (that is *w* when *W* = 1) is complicated by the approach of the system to fixation. When the phenotype frequencies become exceedingly small, dependence of the dynamics of the modifier of *ρ* becomes extremely difficult to detect due to numerical instability; this is especially true for larger values of *k* in *AkBk* regimes when *w* is asmall (Table 1, Figure 6B with *k* ≥ 19).

Figure 6A (see also Table 1) shows that the value 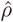 that maximizes the geometric mean fitness is the same as the evolutionarily stable value *ρ*\* in the *A*1*B*1 and *A*2*B*2 selection regimes. For *AkBk* regimes with *k* > 2, our numerical analysis shows the 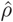 depends strongly on the strength of selection; i.e., the value of *w* relative to *W* = 1. For *AkBl* regimes with *w* = 0.1, the difference between 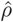 and *ρ*\* is seen even with the *A*1*B*2 environment. For *AkBk* regimes and *w* = 0.9, we find 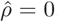, while *ρ*\* is close to 0.9. For larger values of *k*, 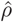 lies between 0.15 and 0.25, while *ρ*\* remains above 0.8 and can reach 0.99 for very large *k*. Comparing Figure 6A with the asymptotic growth rate (AGR) of Xue and Leibler (27), whose parameter *η* is the rate at which an individual learns from its parental lineage, there is a similarity for our curves for *w* = 0.1 and 0.5 with their curve in the *AkBk* environment. They show the AGR decreasing with *η* in the *AkBk* regime for small *k*, but larger values of *k* entail that the AGR has a maximum for an intermediate value of *η*.

Although the models of Xue and Leibler (27) and that analyzed here both incorporate parental and non-parental transmission, they do so in qualitatively different ways. The model treated in the present paper is squarely in the tradition of gene-culture coevolutionary theory, together with modifier theory from population genetics. The different findings from the two classes of models are interesting and suggest that further exploration of the overlaps and discrepancies between the two approaches would be worthwhile.

## ACKNOWLEDGMENTS.

This manuscript was supported in part by the Stanford Center for Computational, Evolutionary and Human Genomics, and by the Morrison Institute for Population and Resources Studies, Stanford University.

